# Optimization of a Translational Murine Model of Closed-head Traumatic Brain Injury

**DOI:** 10.1101/2022.06.02.494542

**Authors:** Brian E. Mace, Eric Lassiter, Evangeline Kamalini Arulraja, Eduardo Chaparro, Viviana Cantillana, Rupali Gupta, Timothy D. Faw, Daniel T. Laskowitz, Brad J Kolls

**Affiliations:** Duke University School of Medicine Department of Neurology Brain Injury Translational Research Laboratory; Duke Health, Department of Surgery, Division of Emergency Medicine; Duke University School of Medicine Department of Neurosurgery; Duke University School of Medicine Department of Orthopaedic Surgery

## Abstract

Traumatic brain injury (TBI) from closed-head trauma is a leading cause of disability with limited effective interventions. Many TBI models impact brain parenchyma directly following craniotomy, and are limited by the fact that these forces do not recapitulate clinically relevant closed head injury. However, applying clinically relevant injury mechanics to an intact skull may lead to variability and as a result, modeling TBI remains a challenge. Moreover, current models often do not explore sex differences in TBI, which is critically important for translation to clinical practice. We systematically investigated sources of variability in a murine model of closed-head TBI and developed a framework to reduce variability across severity and sex. We manipulated pressure, dwell time, and displacement to determine effects on vestibulomotor performance, spatial learning, and neuronal damage in 10-week-old male and female mice. Increasing pressure beyond 70psi had a ceiling effect on cellular and behavioral outcomes, while manipulating dwell time only affected behavioral performance. Increasing displacement precisely graded injury severity in both sexes across all outcomes. Physical signs of trauma occurred more frequently at higher displacements. Females performed worse than males when injured at 2.7mm displacement, and had greater mortality at higher displacements. Stratifying severity based on day-1 rotarod performance retained cellular injury relationships and separated both sexes into injury severity cohorts with distinct behavioral recovery. Utilizing this stratification strategy, within-group rotarod variability over 6 days post-injury was reduced by 50%. These results have important implications for translational research in TBI and provide a framework for using this clinically relevant translational injury model in both male and female mice.

## Introduction

Traumatic brain injury (TBI), especially when caused by closed-head trauma, remains a significant public health concern in both males and females^1^ with mild TBI occurring in up to 3.8 million people annually in the United States alone^2^. In a recent retrospective study of mild traumatic brain injury (TBI), females had a worse outcome than males^3, 4^. These sex related differences in outcome are not explained by sociodemographic or care pathways, and have yet to be identified^3, 5, 6^. Clinically relevant animal models of TBI can provide insight into the mechanisms that underlie the brain response to injury, factors contributing to pathological outcomes, and facilitate screening of therapeutics to improve functional recovery. There are several preclinical TBI models which have recently been reviewed^7–9^. Our model was created in mice as a modification from the Feeney weight-drop model^10^ and leverages a pneumatic piston impact on an intact skull. This approach closely models the biomechanical forces associated with non-penetrating, closed-head TBI, which is the most common form of TBI seen in both civilian and military populations^11–15^. Closed-head injury models have the advantage of replicating clinically relevant injury forces that occur in humans^9^ and have added to the expanding knowledge of the brain response to mild to moderate TBI by increasing the understanding of mechanisms, pathological outcomes, and therapeutic interventions after injury^16–22^. Adapting closed-head TBI models for mice allows researchers to increase throughput and leverage transgenic approaches for mechanistic studies.

One limitation of closed-head TBI models is the potential for relatively high variability in injury severity^23^. In addition, very few studies have systematically evaluated sex differences^7^. The purpose of the current work was to understand the sources of variability in our mouse closed-head TBI model and identify ways to control the level of injury, thus allowing development of a more consistent injury model for both males and females. To explore the basis for the variability, different aspects of the model were systematically explored using assessment of vestibulomotor function and spatial memory in relation to neuronal injury based on fluoro-jade staining in young adult male and female C57BL/6J mice.

## Results

### Baseline rotarod performance was similar between male and female mice

Baseline motor function was assessed via rotarod in 135 male and 144 female uninjured mice. Although female mice had a greater tendency to reach the 300 second time limit than males (27.5% compared to 20%), there was no significant difference in baseline run times between male and female mice (Supplement Figure 1; t-test, p > 0.05).

**Figure 1.**
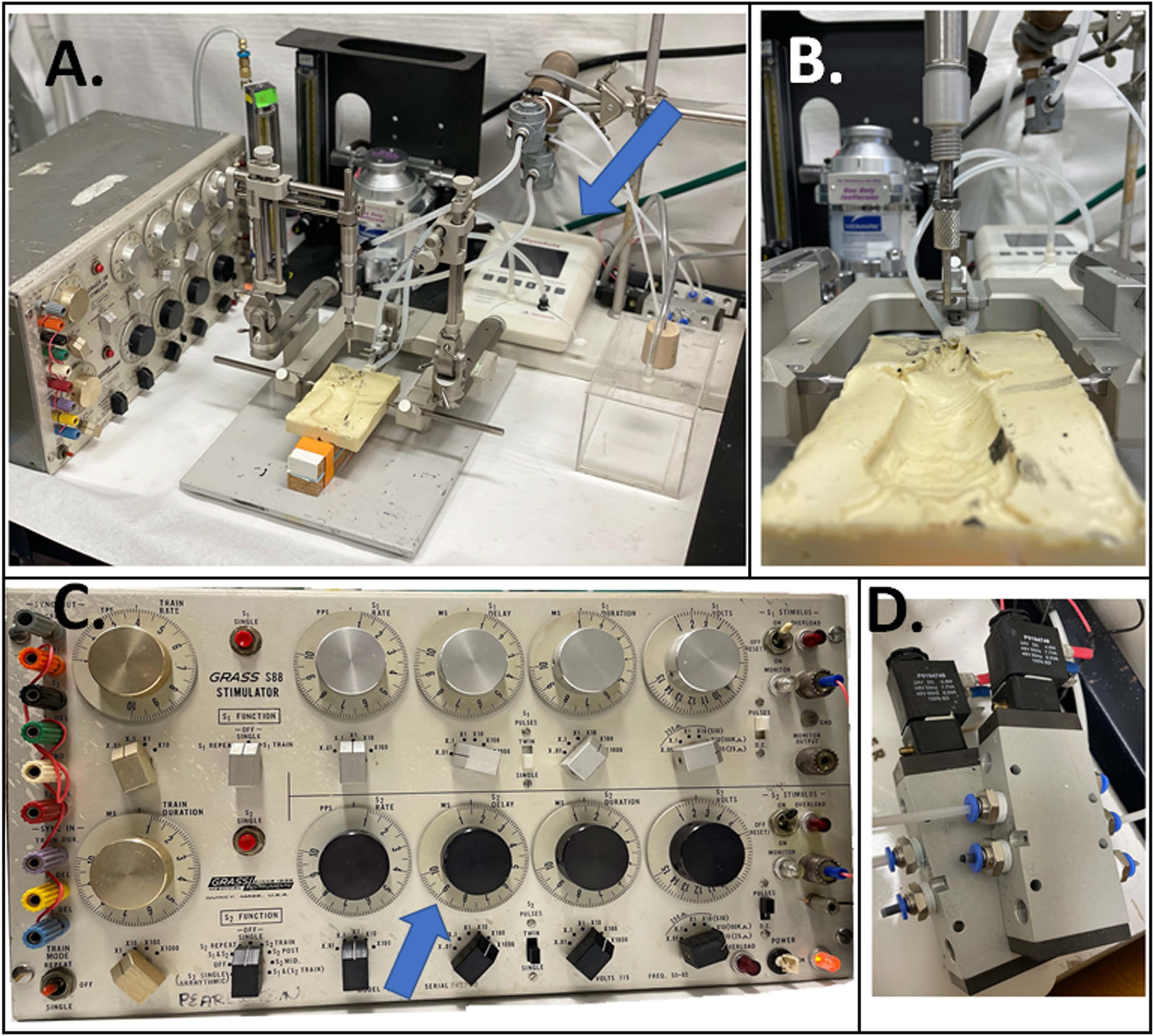
Overview of the pneumatic piston traumatic brain injury induction system (A). Key components include the Grass S88 stimulator (A, C), the solenoid and electronic valves (D) and the ability to control displacement with a stereotactic manipulator. The pneumatic piston (B) is driven by pressure regulated compressed nitrogen. The gas is released in short bursts via solenoid regulated valves (D) driven by a Grass S88 stimulator (C). The piston is driven both down and up by release of a nitrogen gas bolus, requiring two valves. The S88 was wired so that triggering of S1 runs both stimulators (left C). The delay between these pulses is the dwell time for the piston and is controlled by the delay for stimulus 2 on the S88 (blue arrow C). The voltage stimulus output from the Grass is 22-24 V. The anesthesia system is shown by blue arrow in A, and consists of gas mixer (left behind piston) for mixed oxygen and nitrogen supply, isoflurane evaporator (middle A), induction box (right A) and mechanical ventilator next to the evaporator. Note the ear bars of the stereotactic frame are used to hold the plastic mold in place and are not used in the animal’s ears

### Effects of line pressure on injury severity

We employed a translational pneumatic piston impactor model to interrogate the role of device parameters on injury severity (Figure 1). To determine the effect of line pressure on injury severity, male mice (n=10 per group) were randomized to injury using 60, 70, or 90 psi with 150-ms dwell time and 2.5 mm displacement. There were no differences in day-1 rotarod performance between uninjured baseline RR and after injury at 60 psi (one-way ANOVA, F =22.0, p < 0.001, pairwise t-test, p > 0.05; Figure 2A). Higher line pressures of 70 and 90 psi resulted in significantly worse day-1 rotarod performance compared to baseline or 60 psi (one-way ANOVA, F =22.0, p < 0.001, pairwise t-test, p < 0.001; Figure 2A), although no difference was found between 70 and 90 psi (pairwise t-test, p > 0.05). High-speed camera recordings confirmed the piston speed was 33% slower at 60 psi (3.0 m/sec) than at 70 and 90 psi (both 4.5 m/sec).

**Figure 2.**
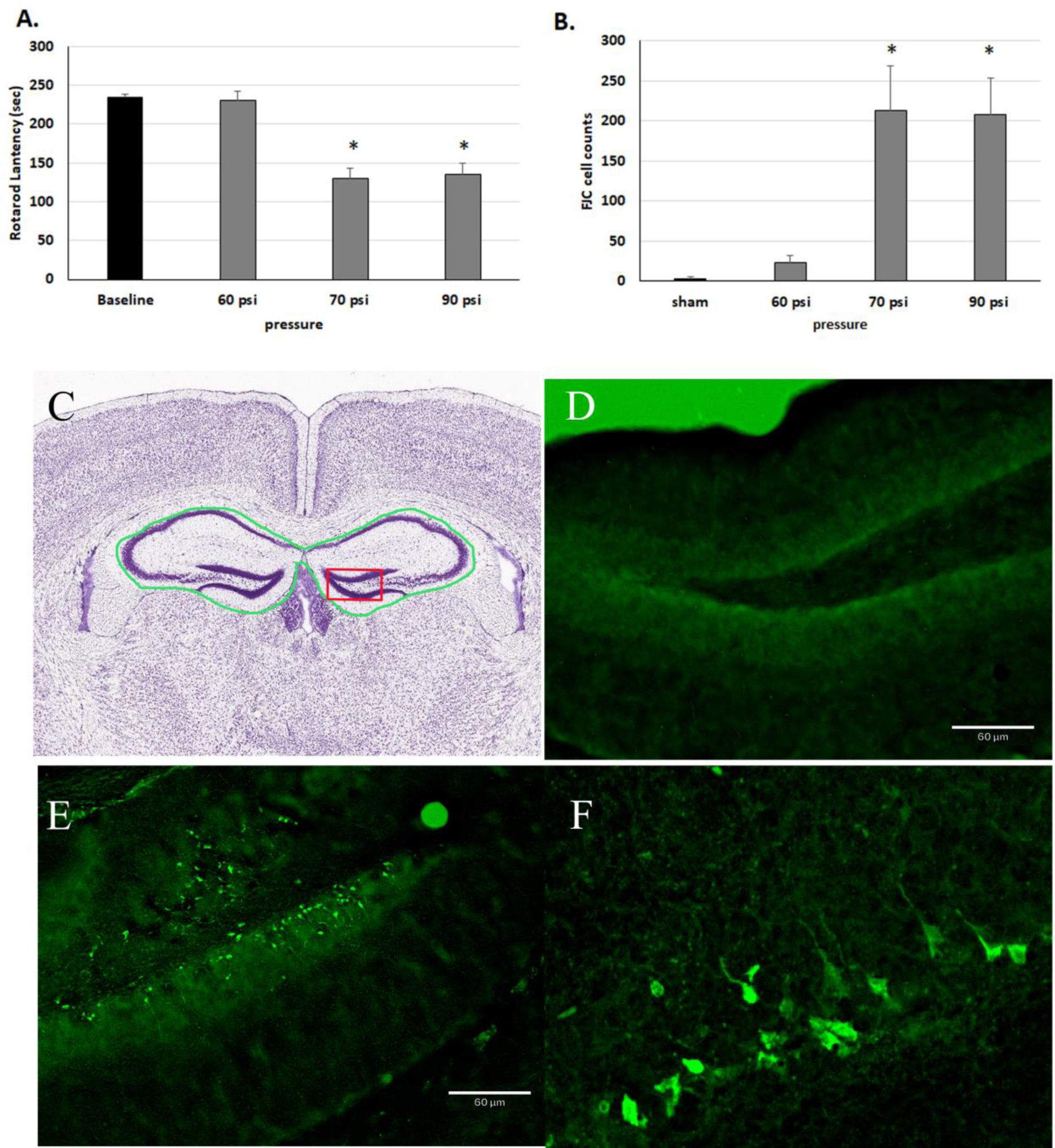
Comparison of injury induced by different piston pressures of 60,70 and 90 pounds per square inch (psi). A) Rotarod performance day-1 after injury was significantly worse (one-way ANOVA, F =22.0, p < 0.001, pairwise t-test, *p < 0.001) for injury induced with pressures of 70 and 90 psi compared to baseline with no difference at 60 psi. B) There is a corresponding increase in the number of fluoro-jade C (FJC) positive cells within the hippocampus of mice injured at the 70-90 psi (t-test, *p < 0.01) compared to sham or 60 psi. Fluoro Jade C (FJC) was used to identify injured/degenerating neurons in the brain after injury. Positive neuronal cells in the first six sections containing the hippocampus bilaterally were counted. C) For reference, a coronal Nissl stain from the Allen Mouse Brain Atlas is shown. The green outline is the area where positive FJC cells were counted and the red box is the approximate location of representative FJC images are shown (D and E). D) A single sham section after injury having no positive FJC staining (20x, bar = 60 um) with-in the dentate gurus. E) FJC positive neurons from a male injured at 3.0mm with FJC positive staining with-in the dentate gurus. F) Digitally magnified capture of FJC neurons identifying injured/degenerating neurons.

Fluoro Jade C (FJC) was used to identify injured/degenerating hippocampal neurons after injury^18^. The total counts of FJC positive neurons in the hippocampus were compared at 24 hours post TBI. The 60 psi impact caused some neuronal damage but this was not statistically different from sham (p > 0.05; Figure 2B). Consistent with day-1 rotarod performance, 70 and 90 psi resulted in substantially higher FJC counts compared to sham (t-test, p < 0.01, Figure 2B). Representative examples of the FJC staining differences are shown in Figure 2 C, D, E and F. Figure 2C, a reference picture of a coronal Nissl stain from the Allen Mouse Brain Atlas^24^. The green outline is the hippocampal areas where positive FJC cells were counted. The red box is approximate location of representative FJC images shown in D and E. Sham animals show little to no positive FJC stained neurons within the dentate gyrus of the hippocampus (Figure 2D, 20x, bar = 60 um) in contrast to the sections from a male mouse injured at 3.0mm, at a PSI of 70 which demonstrated moderate FJC staining within the dentate gyrus of the hippocampus (Figure 2F). Other regions of the hippocampus showed similar differences and total counts within the three main hippocampal regions were used for quantitative comparisons. No differences were found between males and females and so counts were pooled for comparative analyses (p > 0.05). We did not measure any differences in FJC staining between 70 and 90 psi. (Figure 2B). This combined with our high-speed video suggesting no change in the piston velocity suggests there is likely a ceiling effect on injury due to piston pressure increased that is related to the mechanical limitation of the valves used in the nitrogen powered TBI system. Suggesting that calibrating a piston driven model to piston velocity (m/sec) is more relevant than the PSI to trigging the system.

### Effects of dwell time on injury severity

We determined the effects of dwell time by adjusting the S2 delay (Figure 1c blue arrow) to 50-ms (n=9; one animal died and was excluded) or 150-ms (n=10) while maintaining 70 psi and 2.5 mm displacement. There was a graded effect of dwell time on day-1 rotarod performance with 50-ms producing an intermediate deficit (one-way ANOVA, F =16.0, p < 0.001, pairwise t-test, p < 0.01 vs. baseline) and 150-ms yielding the worst performance (one-way ANOVA, F =16.0, p < 0.001, pairwise t-test, p < 0.05 vs 50-ms, Figure 3A). Unlike pressure, increasing dwell time did not affect the amount of neuronal damage by FJC staining (t-test, p > 0.05, Figure 3B). These results suggest that, unlike varying piston velocity, modifying dwell time may not provide a practical way to vary injury severity within the model as we noted a dissociation between behavioral endpoint measures and underlying cellular injury.

**Figure 3.**
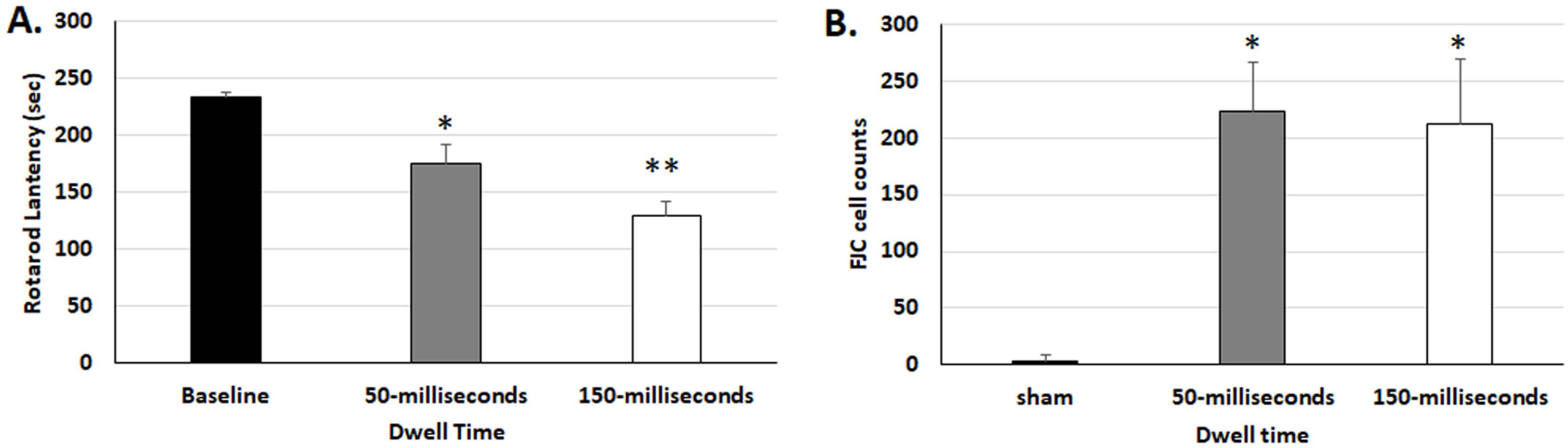
Comparison of injury induced by different piston dwell times. A) Rotarod performance day-1 after injury was significantly worse (one-way ANOVA, F =16.0, p < 0.001, pairwise t-test, *p < 0.01) for injury induced with dwell times of 50 milliseconds (ms) compared to baseline. The day-1 latency for the 150-ms dwell time was significantly different from both baseline and the 50-ms dwell times (*p < 0.05 and *p <0.01, one-way ANOVA pairwise t-test). B) There is a corresponding increase in the number of fluoro-jade C (FJC) positive cells within the hippocampus for both dwell times in injured mice, (t-test, *p < 0.01,), however, no difference was observed between 50 and 150-ms FJC counts (t-test, p > 0.05,) 1 day after injury.

### Effects of head displacement on injury severity

The final variable considered was the extent of head displacement by the piston. A total of 279 mice (144 females and 135 males) were randomized to receive 2.0, 2.3, 2.5, 2.7, 3.0 or sham (0.0 mm) displacement with 70 psi and 150-ms dwell time. Post-injury rotarod and Morris Water Maze (MWM) performance were compared between cohorts. The aggregate results of varying displacement in all 279 mice are shown in Figure 4A. As head displacement is increased there is a corresponding increase in functional deficit, evidenced by shorter rotarod latencies. At displacements of 2.5 mm and above, some mice fail to fully recover back to baseline rotarod performance by 6-days post-injury (baseline vs. day 6 RR times: Sham-2.3mm, t-test p>.05, 2.5-3.0mm p<.0.01). Increased displacement and corresponding decline in day-1 rotarod performance is associated with increased neuronal damage measured by FJC positive cells (multiple linear regressions, R^2^ = 0.997, p < .05, intercept at p <0.05) (Figure 4B).

**Figure 4.**
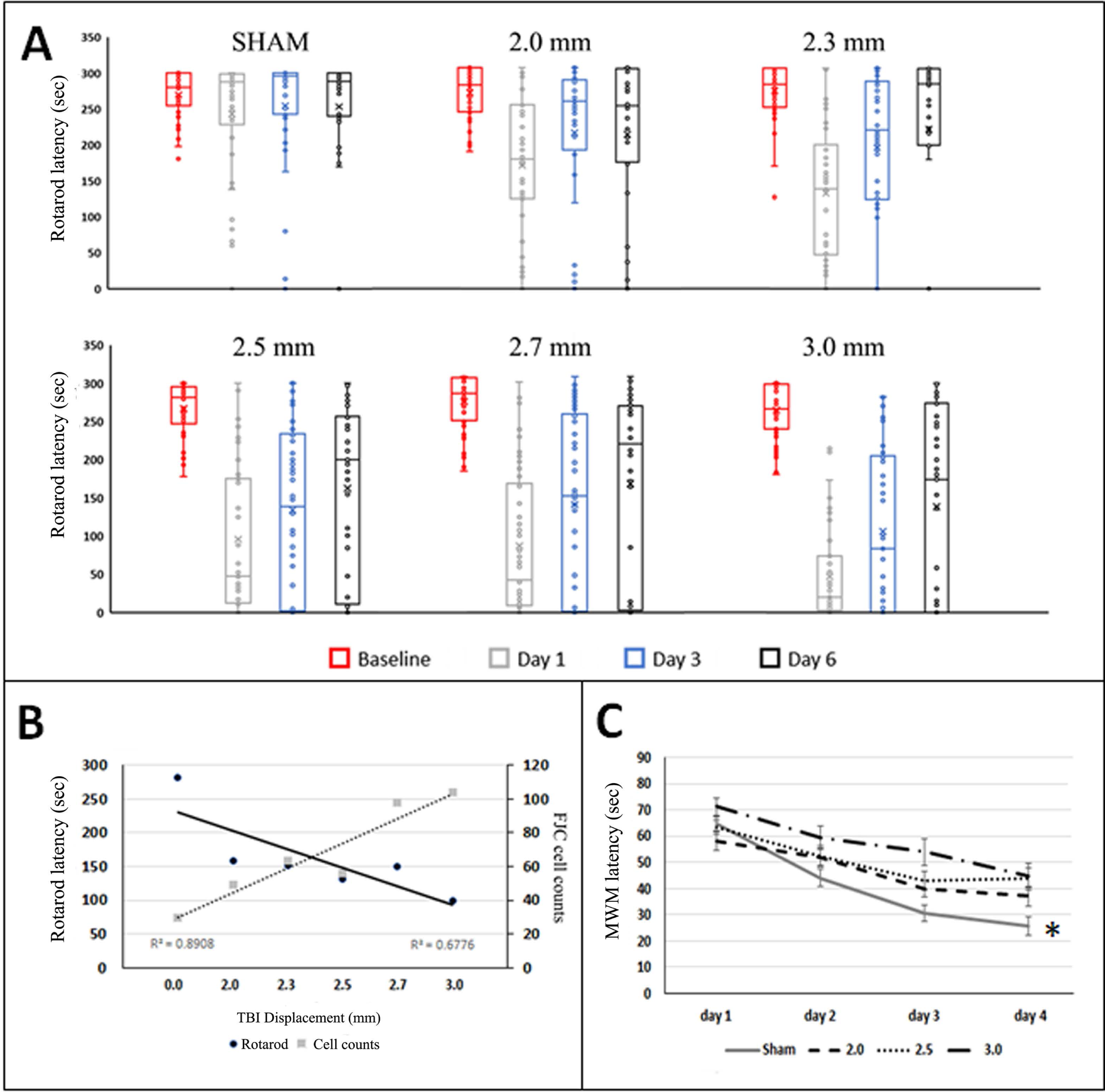
Aggregate rotarod latency performance after injury from both male and female mice at increasing displacements. A) As displacement increases by small (0.2-.03 mm) amounts, the latency for falling off the rotarod task one day after injury declines (light gray boxes). The animals have a characteristic recovery back to baseline latencies by day 6 post injury (black boxes). Note some mice do not return to baseline with displacements of 2.5mm or greater. B) Fluoro Jade C (FJC) positive cell counts increase with increasing displacement, consistent with the decrease in rotarod latency times post injury. C). Morris Water maze (MWM) performed at 21-28 days after injury demonstrates significant (ANOVA, *p < 0.05) increased latency to finding the platform with increasing displacement for all displacements compared to sham (2.3 mm and 2.7 mm not shown).

Similar effects were observed in MWM performance. As displacement increased, performance worsened with longer times required to find the platform. Learning curves from injured mice were significantly worse than sham-injured animals, even at the smallest displacement (2.0 mm; two-way ANOVA, F =5.29, p < 0.05, Figure 4C). Of the tested parameters, these data suggest that the most salient and practical variable to modify injury severity is head displacement

### Observed physical injury and behavioral effects of acute trauma

Physical signs of trauma included skull fractures, skull hemorrhages, and intracranial hemorrhages. These signs more frequently occurred at larger displacements across all animals (n=240; Figure 5A), and no sex differences were seen (p >0.05, data not shown). Over 75% of mice with 2.3 mm displacement or less showed no fractures or had small, linear fractures with approximately 24% having a fracture with some hemorrhage (Figure 5A). At larger displacements (2.5 and above), 85% of mice had fractures and 62% had fracture-associated hemorrhages. Displacement of 2.7 and 3.0 mm resulted in occasional compound fracture that deformed the skull, depressing parietal bones and compressing the cortex. On several occasions, these open fractures were associated with brain extrusion. With displacements between 2.5 to 3.0 mm, intracranial hemorrhages were occasionally noted and were associated with increased mortality.

**Figure 5.**
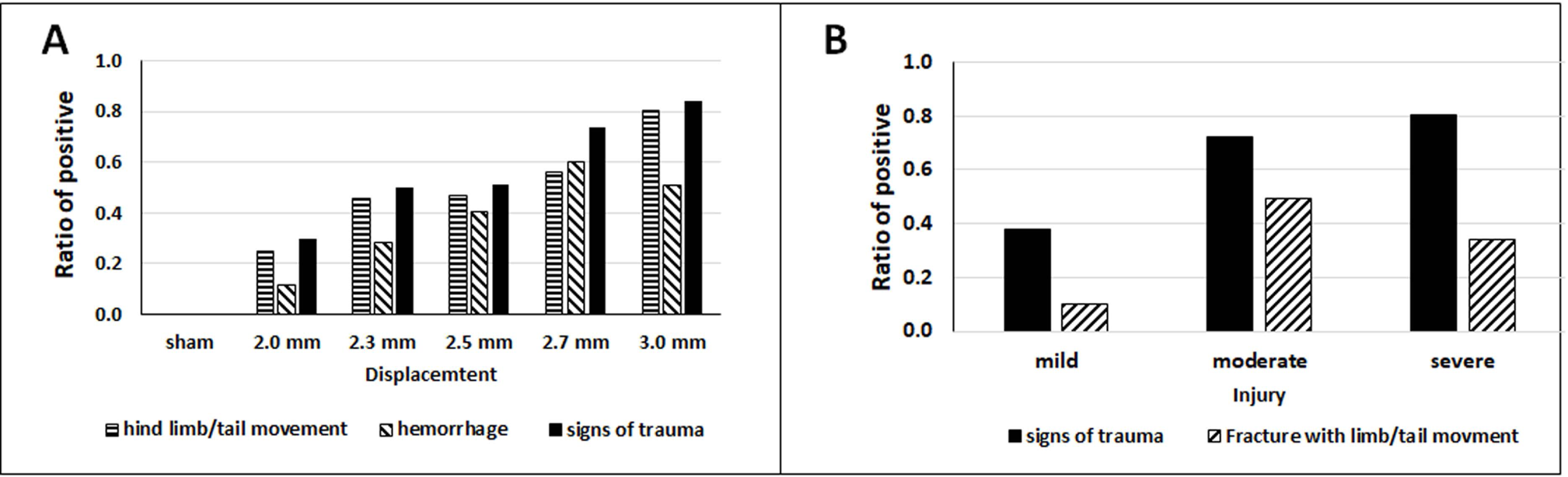
Clinical signs of injury are more commonly seen with greater injury severity settings. A) Twitching and jerking of the hind limbs and Straub tail were more frequently seen in association with higher displacements. Fractures of the skull and hemorrhage were also more common at higher displacements. B) Grouping the animals based on day 1 rotarod performance which more closely reflects actual injury severity, accentuates these clinical associations with injury. These results suggest these clinical signs of injury can be a reasonable surrogate for adequate delivery of TBI injury.

A subset of animals had clinically observed tonic and/or clonic movements of hind limbs and tail raise (i.e. Straub tail^25^) that occurred either independently or simultaneously between 2- and 5-seconds post-impact. As with skull fractures, this phenomenon occurred more frequently with larger displacements, with up to 80% of mice with 3 mm displacement demonstrating these signs, compared to ∼45% or less with displacements <2.5 mm (Figure 5).

These physical manifestations of injury had no statistically significant association with day-1 rotarod performance within a given displacement cohort (t-test, p>0.05). However, approximately 75% of all injured animals experienced one or more of these physical manifestations (n=180), with 21% of the animals having all three observations (n = 50), 32% have at least two different observations (n=76), 23% having at least one (n=54), and 24% having no physiological manifestations at all (n=58) (Figure 5A)

When assessing for conclusively severely injured animals (displacement 2.7-3.0 and Day 1 RR < 50 seconds, n=72) less than 10% of the animals showed no sign of trauma, with 30% (2 out of 7) of those animals having full RR recovery by day 6, suggesting a more mild injury in these mice (data not shown). These results support the potential use of physical signs of trauma as an indicator of more severe injury when higher displacements are used.

### Female mice reach maximal injury at smaller displacements

No sex differences occurred at displacements less than 2.7 mm (2.0 and 2.3 mm not shown, two-way ANOVA p>0.05). However, at a displacement threshold of 2.7 mm, females had significantly lower rotarod latencies post-injury (two-way ANOVA, F=33.5, p < 0.001; Figure 6A). Both male and female mice demonstrated poor rotarod performance at 3.0 mm displacement and were not significantly different from each other (Figure 6A). There were no sex differences in spatial learning by MWM across tested displacements (data not shown). These data demonstrate that displacement can be used to scale injury severity in both male and female mice and that females reach an injury ceiling at 2.7mm of displacement.

**Figure 6.**
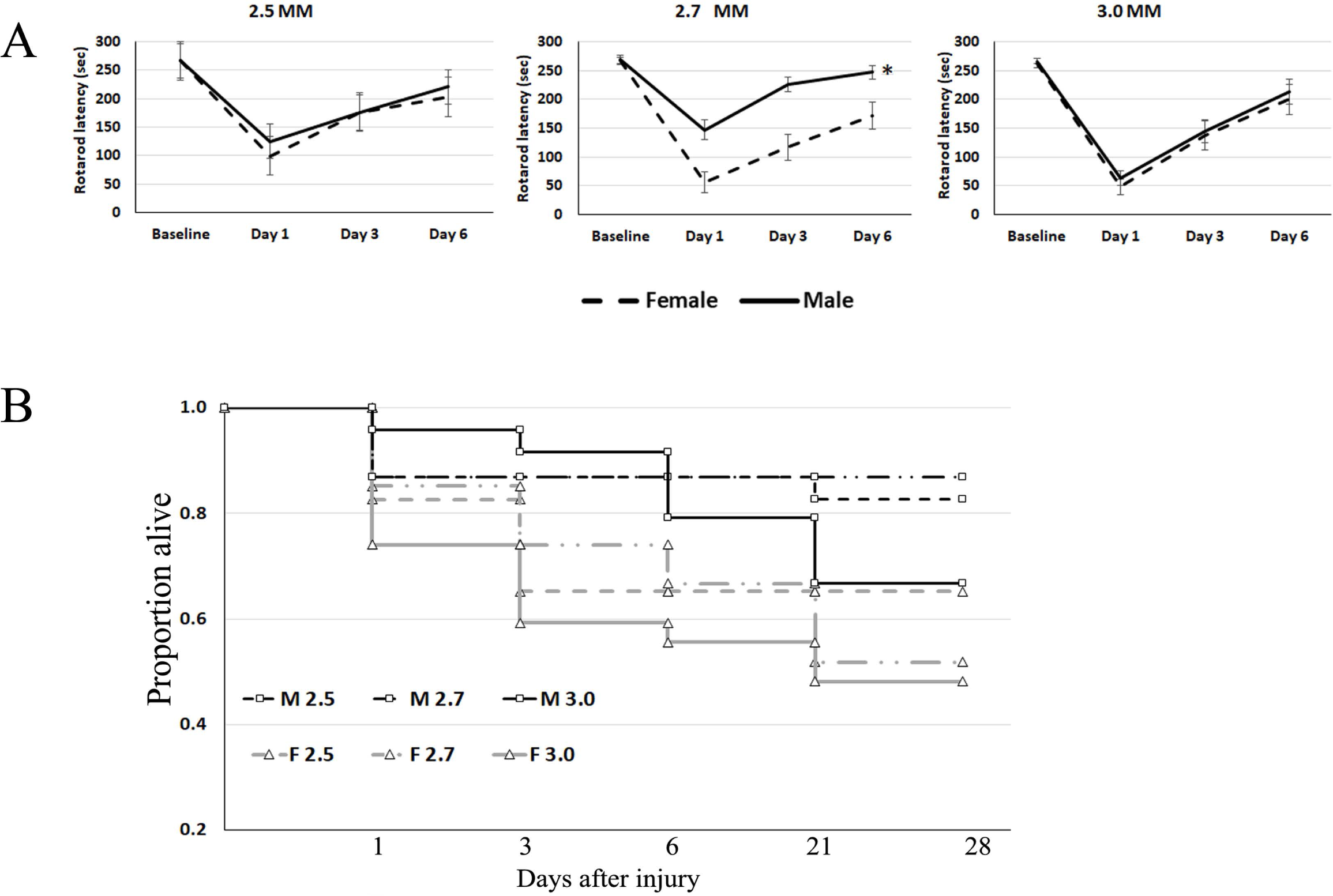
A) Rotarod data of male and female mice following larger head displacements. Left panel, rotarod latency with a displacement of 2.5 mm in female and male animals were not significantly different (p > 0.05). Middle panel demonstrates that females compared to males had significantly lower post-injury RR latencies at a displacement of 2.7 mm (ANOVA, F=33.5, *p <0.0001). However, the female latencies at 2.7mm were not different from the male latencies at 3.0mm (ANOVA, p > 0.05). Right panel shows that at the maximum displacement tested (3.0 mm) both male and female animals have similar RR recovery curves with no significant differences (p > 0.05), though mortality was higher in females at these larger displacements. B) Kaplan-Meier post injury survival curves for males and females with higher injury severity. At lower displacements of injury, survival was similar, however as the displacements increased from 2.5-3.0 mm mortality was significantly higher in the animals (p < 0.001, log-rank test). At displacements of 2.7 and 3.0 mm, females had a 28-day survival of approximately 50%.

### Model associated survival

For all animals (n=279), this model had a 6-day survival rate of approximately 80% across all displacements (end of rotarod testing) and final survivability of ∼80% at 35 days (end of MWM testing). Male mice had a 90% survival rate while females had a survival rate of 70%. Not surprisingly, 70% of censored animals (n=57) came from larger displacements (2.5 mm and above; Figure 6B). Female mice accounted for 75% of all deaths (n=43), with ∼70% coming from the higher displacements. Just over half of the female mice (56%) survived the 3.0 mm displacement, in contrast to 80% of males surviving this degree of displacement (Figure 6B). Kaplan-Meier survival curves for males and females across each of the higher displacements are shown in Figure 6B. Log-rank test on 2.5, 2.7, and 3.0 mm displacement groups demonstrated significant differences between males and females in both cohorts of pooled injury (p < 0.001; lower displacements not shown).

### Optimizing the model for male and female injury comparisons

Higher mortality in females suggested that smaller displacements would be needed to induce similar survivable injuries compared to males. Given our observed relationship between day-1 rotarod performance and neuronal damage, we considered day-1 rotarod performance as a surrogate to stratify injury severity. Based on day-1 rotarod performance, females at 2.7 mm displacement had similar injury severity as males at 3.0 mm displacement (Figure 7).

**Figure 7:**
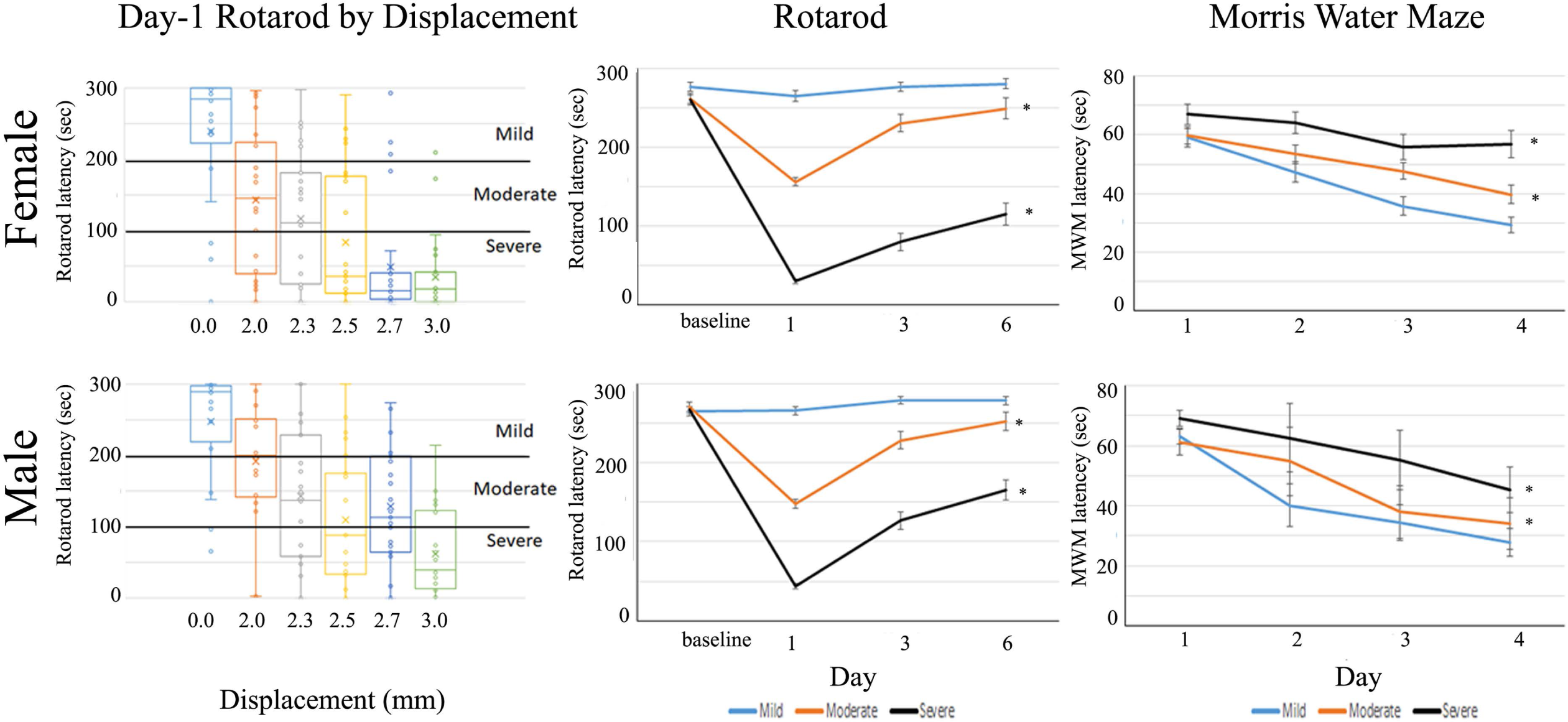
Grouping animals by rotarod performance one day post injury lowers variability and identifies three distinct injury groups. Female (top row) and male (bottom row) C57Bl/6 mice were systematically injured with incrementally greater head displacements (0, 2.0, 2.3, 2.5, 2.7, and 3.0 mm, n=20 per group for each sex). The box whisker plots on the left demonstrate the effects on rotarod testing one day after injury. As head displacement increases, the latency time to falling off the rotarod decreases indicative of worse injury. Mice were placed into three categories based on day-1 rotarod performance (left panel categories), mild, moderate and severe. (middle panel). When grouped by day-1 injury severity, subsequent performance on rotarod on day 3 and 6 is significantly worse as a function of injury severity (ANOVA, *p < 0.0001 for both males and females). Similarly, when mice were tested in Morris Water Maze (WM) (right panel) outcomes could be categorized into: good (< 40 sec), intermediate (40-60 sec), and poor (> 60 sec) and corresponded with the rotarod day-1 grouping (ANOVA, *p < 0.0001 for both males and females).

Mice with day-1 rotarod latencies greater than 200 seconds were considered to have mild injury, as they were associated minimal neuronal damage, and short recovery times with full recovery by day 3 post injury. Rotarod latencies between 100 and 200 seconds were considered moderate given intermediate neuronal damage, rotarod recovery to baseline by day 6, and observed behavioral signs of injury associated with this group. Given high neuronal damage, mortality, and incomplete motor recovery in mice with day-1 rotarod latencies less than 100 seconds, we consider this to represent severe injury (Figure 7). Mice with day-1 rotarod latencies of 10 seconds or less (n=46) had a 78% mortality rate, with a >50% survival rate at a latency of ∼ 20 seconds.

Grouping mice by day-1 rotarod performance retained the relationships to underlying cellular injury in FJC staining (Figure 8) and clearly separated mice of both sexes into distinct cohorts with significantly different rotarod recovery curves (two-way ANOVA, Female, F= 286.5, p < 0.001; Male, F=260.3, p <0.001; Figure 7) and MWM performance (two-way ANOVA, Female F=15.6, p< 0.001; Male, F=27.4, p < 0.001; Figure 7). Severely injured mice did not fully recover rotarod performance by day 6 (t-test, both sexes p < 0.001) and showed impaired spatial learning in the MWM task (Figure 7), (two-way ANOVA, both sexes p < 0.001).

**Figure 8.**
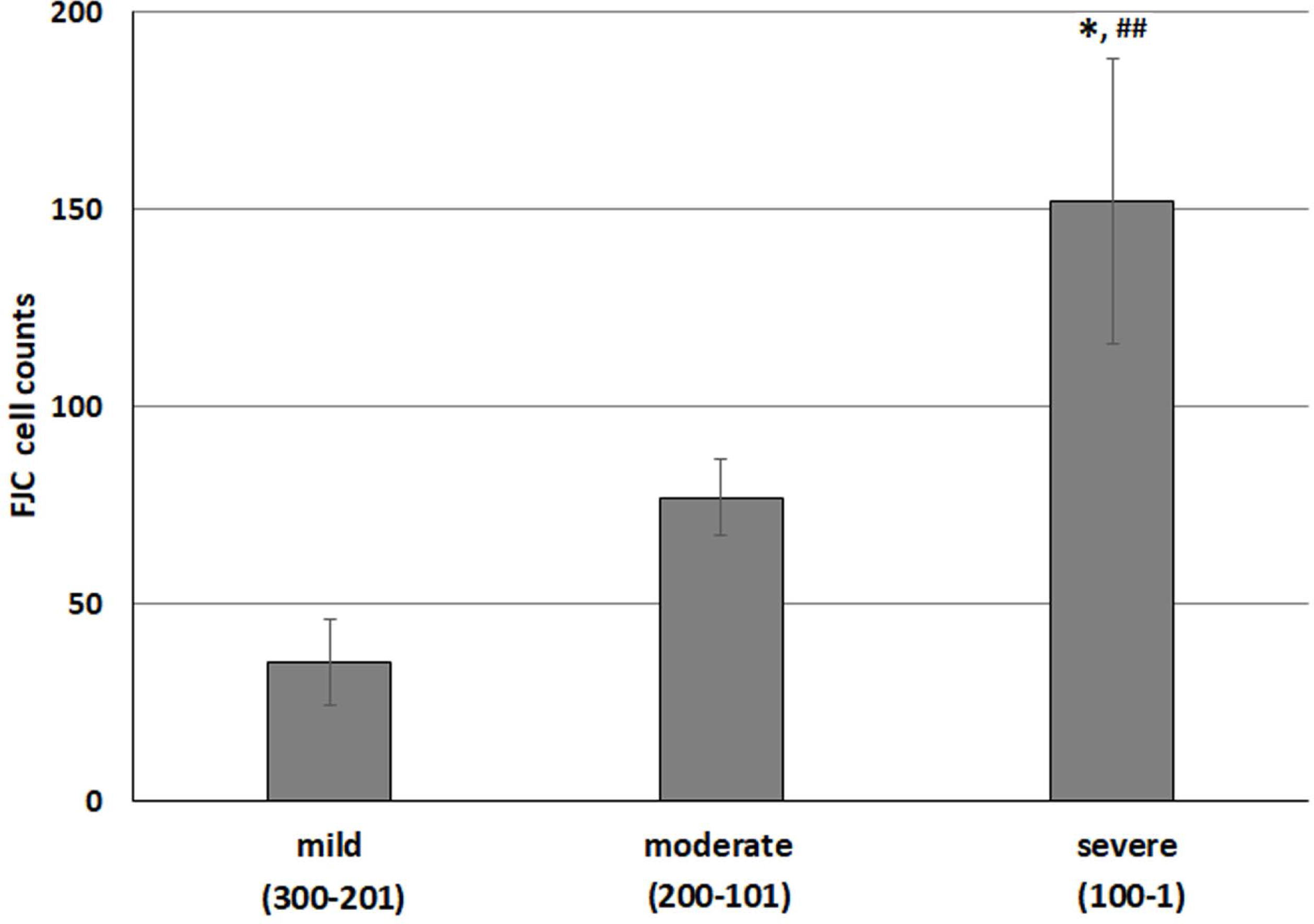
FJC staining is increased in the mice grouped by RR day one latency performance; mild (201-300 seconds), moderate (101-200 seconds) and severe (≤100 seconds) instead of displacement. FJC positive counts significantly increase with more severe TBI from mild to moderate/severe (t-test, *p ≤ 0.01) and from moderate to severe (t-test, ^##^ p < 0.05). All are significantly elevated compared to sham (p > 0.05)

We also grouped mice by displacement into mild (0.0-2.0 mm), moderate (2.3-2.5 mm), and severe (2.7-3.0) cohorts and directly compared values to stratifications based on day-1 rotarod. Although Figure 7 shows that overall outcomes between the two are similar, there were statistical differences that were observed between approaches (supplemental table 1). Stratification by day-1 rotarod performance reduced overall rotarod latency variability over the 6 days of testing by approximately 50% (supplemental table 1). Additionally, 75-80% of the animals in the moderate to severe stratifications had at least one sign of injury (fracture or abnormal motor activity, Figure 5B). The presence of these observations had a positive predictive value of 0.78 and their absence has a negative predictive value of 0.90 for day-1 rotarod performance less than 100 seconds.

## Discussion

The purpose of the current work was to understand the sources of variability, define injury severity cohorts with highly similar histologic markers of injury severity, and identify any sex differences in our well-established pneumatic piston model of closed-head TBI. We found that pressure and dwell time do not provide a practical way to scale injury, and these parameters can be set to minimize their contribution to variation in injury outcomes. In contrast, head displacement was highly scalable and allowed for identification of an injury severity in females that closely approximates the mortality and behavioral outcomes seen in male mice. Our results have important implications for translational research in TBI and provide a framework for using this clinically relevant injury model in both male and female mice.

We explored three sources of variation inherent with the pneumatic piston driven closed-head TBI model, namely piston line pressure, dwell time, and displacement distance. Although both pressure and dwell time had clear effects on both cellular and behavioral outcomes after injury, we found the effects were not scalable in a practical way. In contrast, displacement provided exquisite control of injury severity in both males and females. Displacements varied by 0.2 - 0.3 mm showed consistent, reproducible, and statistically significant differences in the resulting motor function, spatial learning and memory, and direct histological damage within the hippocampi of injured animals. In general, female mice were more susceptible to larger displacements with worse rotarod performance and lower survival at larger displacements compared to the male mice. Based on the systematic exploration of our three parameters we conclude that optimal injury in terms of highest survival and maximum severity is obtained using 70 psi with a velocity of 4.5 m/sec, 150-ms dwell time, at a displacement distance of 3.0 for males and 2.7 mm for females. This study was limited to only animals that survived injury and to a single genotype background, both of which may change these parameters and outcome depending on experimental design.

Although we automated the model by using a GRASS S88, which reduced variability in dwell time, a high degree of variability is common with model development, and supervised training by experienced personnel is required for model consistency. The sources for this variation are not clear but are likely related to procedural factors, such as handling between cage and injury system, intubation skills, and speed of injury procedural performance which varies the anesthesia duration. The current study used the same, highly experienced surgeon (author EC) for all procedural injuries so these factors were controlled in the current work.

While displacement provided a practical method for controlling the severity of injury, it still resulted in variable injury levels within a given displacement group, particularly at displacements greater than 2.5 mm. The variability could be significantly reduced by using day-1 rotarod latency as a behavioral surrogate for injury severity. We demonstrated that by grouping mice into mild (200 seconds or more), moderate (100-200 seconds), and severe (< 100 seconds) cohorts based on rotarod performance, the variability between mice within a given cohort was reduced by 40-60% and was closely associated with the degree of neuronal damage as measured by FJC (Figure 8). We also noted that a subclass of the severely injured mice, those that ran for less than 10 seconds on rotarod had a nearly 80% mortality which allows for early removal of these animals and further contributes to producing more homogeneous cohorts of animals.

In the situation where severe injury is intended, the absence of any physical sings of injury indicate there is less than a 10% chance that the animal is severely injured and the animal could be removed from the cohort to improve injury homogeneity. The reduced within-cohort variation greatly improves the power of the model.

Our stratification approach using rotarod performance on post injury day 1 reduced the variation in mean rotarod times within a cohort to 10% or less. This results in 90% power to detect a 15% difference in the means with an alpha of 0.05 and beta of 0.1 with just 9 mice per arm. This dramatic reduction in the number of animals needed further increases the throughput and value of the model. Stratification by rotarod performance also allowed for injury equivalence between males and females despite needing to use less displacement in the females. Our proposed clinical stratification of the animals is similar to one used in human TBI patients to assess injury severity. In clinical practice, the Glasgow Coma Scale (GCS) is an evaluation of exam performance that includes motor responses, eye opening and verbal responses to stimulation^26^. Patients with TBI are stratified into mild (GCS 13-15), moderate (GCS 9-13) and severe (3-8) injury based on the score. While relatively crude, this tool is predictive of outcome for moderate to severe injury patients^27–29^. This lends further validating support to our model which not only employs clinically relevant injury mechanisms, but also demonstrates close associations between clinical assessment, underlying cellular injury, and mortality risk.

Our model has been an effective tool in the field of TBI for many years^16, 18, 22, 30^, and has the advantage of being relatively high-throughput with the capacity to perform as many as 30 consistent, reproducible, TBI injuries per day. By virtue of being in mice, this model allows for the use of transgenic approaches to study specific roles of various pathways and mediators of TBI response by the brain, and our proposed approach to cohort selection based on rotarod performance can further optimize throughput for both males and females. However, the model still has some important limitations. In our experience, day-1 rotarod performance should be assessed 16-24 hours (next day) after injury as the stress can increase mortality in more severely injured animals when run at earlier times. Thus, treatments within the first 24 hours post-injury may alter day-1 rotarod performance and preclude its use as a stratification tool, this method should not be used to look at genetic differences, or used to test any variability that is expected to be elevated within the first 24 hours prior to stratification. Nonetheless, we did find that some signs of injury, such as skull fracture, hind limb twitching, and acute recovery times may provide alternative surrogates for injury severity. Our results suggest that in an experiment in which significant injury is intended, animals which lack these clinical signs at the time of injury can be removed to reduce the variability and provide a more homogeneous injury condition for comparing outcomes. When combined with mortality, removal of these animals will require ∼20% more animals than the number calculated as needed per arm of the study but will have less variability in later behavior and histological outcomes, improving rigor and reproducibility.

This model leverages clinically relevant injury forces common to both military and civilian populations^31–34^. The inclusion of females will allow greater pre-clinical testing in both sexes prior to human translation and will facilitate the exploration of sex differences in the response to TBI. We have now further optimized our high throughput model by demonstrating scalable injury severity using clinically relevant forces and producing easily measurable outcomes that mimic human clinical injury severity assessment (GCS) to functional outcome and mortality in both males and females. This will facilitate translational research in TBI and provide a framework for potential biomarkers, transgenic exploration of key injury response mediators of inflammation, therapies and their mechanisms, and define pathophysiological differences between severe and mild TBI as well as their interaction with sex in a pre-clinical model.

## Methods

### Animals

All Animal procedures were approved by Duke University’s Institutional Animal Care and Use Committee. Duke University maintains an animal program that is registered with the USDA, assured through the NIH/PHS, and accredited with AAALAC, International. Recommended ARRIVE guidelines were used in design of all animal research. Male and female C57BL/6J mice (Jackson Laboratories, Bar Harbor, ME) were purchased at 8-weeks of age and allowed to acclimate for at least 10 days prior to use. Animals were housed in 12-hour day/night light cycle with continuous access to food and water.

### Randomization and blinding

Animals were randomly assigned to groups prior to surgery using a block randomization scheme to ensure that parameters were explored systematically over time, across groups, and between sexes as we were limited to 20 injuries per day. There were six different graded injury severity paradigms (characterized by increasing head displacement sham, 2.0, 2.3, 2.5, 2.7, 3.0). These were broken down into twelve different surgical injury days (12 blocks), each block containing five displacements utilizing twenty mice 10 of each sex (example: block 1 [sham,2.0,2.3,2.5,2.7,] block 2 [2.0,2.3,2.5,2.7,3.0] etc.). The order of the twelve blocks was randomized by utilizing a random number generator (Excel 2019, Microsoft). Each cohort of animals was then randomly assigned to the specific injury displacement parameters from within the block on the day of injury. All animals were used for the Kaplan-Meier curves but, only surviving animals were tested on rotarod on days 1, 3, and 6 after injury and then subsequently tested in Morris water maze at day 21-28 post-injury, unless assigned to an early histologic endpoint. Animals that died prior to reaching the assigned endpoint had their data censored, and their injury parameter and endpoint was replaced and reassigned to a future cohort. Investigators associated with histological processing and behavioral assessments were blinded to the injury assignments.

### Closed Head Injury Model (CHI)

The murine closed-head TBI model used here was modified from rats^35^ and is previously described^16, 17, 22^. To eliminate the interrater variability of our previous manual valve system, we modified our system to allow full automation of the impact. Adjustable variables include: the nitrogen gas line pressure in pounds per square inch (psi) that is measured after the regulator, this extends (down) and retracts (up) the piston; the time the piston remains fully extended (dwell time); and the distance the head is displaced by the piston upon impact (displacement).

### Anesthesia and Surgery

10-12 weeks old mice were anesthetized with 4% isoflurane (Patterson Veterinary, Greeley, CO) followed by oral endotracheal intubation. Mechanical ventilation (PhysioSuite, Kent Scientific, Torrington, CT) occurred at a 0.3 mL tidal volume with 30% oxygen and 70% nitrogen mixed with a maintenance dose of isoflurane (1.6%). Using a RightTemp (Kent Scientific, Torrington, CT) thermal regulation system, animal body temperature was maintained at 37° C using heat lamps. Animals were placed in a plastic mold and positioned within a stereotactic frame. To avoid basilar skull fracture, the ear bars were not secured and the head was not fixed within the frame. The skull was exposed via midline incision in the scalp and a concave metal disk (3 mm diameter) was secured to the skull ∼1 mm caudal to bregma. Isoflurane was discontinued just prior to impact. Following impact, the incision was closed with staples. Animals remained intubated and were extubated once they recovered spontaneous ventilation. Following extubation, mice were monitored under a heat lamp until mobile and then were transferred to a new cage with free access to food and water. Sham animals went through all procedures except contact from the piston, these animals had a slight (mild) injury when compared to animals that did not go through the procedure, and are included in mild injury cohort.

### Pressure Regulation

Animals were injured using a pneumatic impactor with a 2 mm ridged piston (ARO Silverair SD05-P4B4-004 C9, Air-Power, Inc High Point, NC) driven by non-flammable, compressed nitrogen. The piston delivered a single midline impact to the disk affixed to the skull, which distributed the force of the impactor, thus reducing likelihood of fracture and mortality. Piston speeds and dwell times were confirmed by analyzing video of piston movement recorded at 1,000 frames per second via high-speed camera (Sony RX10 IV, Model No: DSCRX10M4/B).

### Control of the Piston with Electronic Valves

Triggering of the pneumatic impactor was controlled using a GRASS S88 Stimulator (GRASS Instruments, Quincy, MA, Figure 1A, C) which allowed for precise control over dwell times between 50- and 150-milliseconds (ms). The GRASS S88 was wired to function on a single-button press using the stimulator 1 (S1) button to trigger two output pulses, one from S1 and one from stimulator 2 (S2), at an adjustable interval delay (Figure 1C left). Stimulator outputs would trigger the two solenoids (Parker, P0194749, Richland, MI) controlling gas flow through air piloted valves (Parker, P2LBX592ESHDDB49, Richland, MI) to release pressurized nitrogen gas which initiated extension and retraction of the piston, respectively (Figure 1B, D). Piston dwell time was controlled by using the S2 delay (Figure 1C blue arrow) set to the desired time. To avoid repeat trigging, single pulse was selected for both S1 and S2 and the train rate was set to zero. An output voltage of 20-24V was required to trigger the solenoid valve without error.

### Control of Displacement

Impact displacement was varied by using a stereotaxic micromanipulator (0.1 mm resolution; Stoelting, Wood Dale, IL) to position the piston over the head (Figure 1B). Positioning occurred by first fully extending the piston manually and lowering it via the micromanipulator until it contacted the metal disk on the skull. The piston was then manually returned to its fully retracted position. The impactor was then lowered by the desired displacement using the micromanipulator. Selected displacements included 0.0 (sham), 2.0, 2.3, 2.5, 2.7, and 3.0 mm. While ear bars were not used, the endotracheal tube was taped in place along with the tip of the nose to prevent any head movement prior to triggering the impactor. A fixed, intermediate displacement of 2.5 mm was chosen for pressure and dwell time experiments to allow for more or less injury to be induced by the parameter changes.

### Behavioral Testing

All behavioral testing occurred in a room permanently assigned for behavioral testing. Vestibulomotor function was assessed using an automated Rotarod ENV-577M (Med Associates Inc., Georgia, VT)^17, 18, 22^. Mice were trained for three days prior to injury. Training consisted of four trials with a minimum of 15 minutes between trials. For the first training trial, mice ran at the initial speed of 2 revolutions per minute (rpm) accelerating to 20 rpm over 300 seconds. Mice who stopped walking or fell prior to the 300 second endpoint were gently placed back onto the rotarod. This procedure was repeated for the remaining three trials but with speeds of 4 rpm accelerating to 40 rpm over 300 seconds. Baseline times were collected 1 day prior to injury on the accelerating 4-40 rpm setting with the mean time of the three trials being used as baseline performance. Trials were stopped when mice either fell off the rotarod or looped three consecutive times without walking. On subsequent days 1, 3 and 6 after injury, animals were tested 3 times on the 4-40 rpm accelerating setting with at least 15 minutes of rest between runs. Reported results were the mean of the three trials for that day for each mouse.

The Morris water maze (MWM) is a well-established assay of spatial learning^36, 37^ and tests the animal’s ability to find a 7.5-centimeter diameter hidden platform in a 120 cm diameter pool within a 90 sec interval using visual cues. The pool was filled the day before testing to allow the water to acclimate to room temperature (24±1 °C). Latency times to find the platform were recorded from above and analyzed using Anymaze (V 6.3.4 Stoelting, Wood Dale, IL). On the first testing day, mice were placed on the platform to allow tank observation and orientation to visual cues without the stress of being in water. Mice typically jumped off the platform after a few seconds, and swimming ability was observed up to 30 seconds to confirm testing capacity before being returned to the platform. During testing, mice were placed at the edge of one of four quadrants for each of four trials. If unable to find the platform after 90 seconds, mice were assisted to the platform and left there for 10 seconds to orient using spatial cues. Following removal from the pool, mice were placed in a cage warmed by a heating lamp (air temperature 35-37° C) where they remained for an inter-trial interval of at least 15 minutes. Four trials were conducted on four consecutive days and the mean time was used for each day’s performance measure.

### Fluoro Jade C

Fluoro Jade C (FJC; Histochem, Jefferson, AR) was used to identify injured/degenerating neurons in the brain after injury^38^. Animals were euthanized via isoflurane inhalation and transcardially perfused with 0.9% saline. Brains were carefully dissected before immersion fixation for 16-24 hours in 10% neutral buffered formalin (VWR, Radnor, PA) and cryoprotection in 30% sucrose for 24 hours. Whole brain coronal sections (40 μm) were cut on a sliding microtome and collected in cryoprotectant solution containing ethylene glycol, sucrose, and sodium phosphate. Every twelfth section, covering 480 μm between sections, was mounted onto a gel-subbed charged slide and used for staining. Briefly, after mounting and drying overnight, slides were immersed in 1.0% sodium hydroxide, 80% ethanol for seven minutes and washed in deionized water. Slides were incubated in potassium permanganate for 10 minutes to reduce background staining, then washed and incubated in the dark in 0.001% FJC for 10 minutes. Slides were washed again in deionized water, then dehydrated and cover-slipped with DPX mounting medium (Fluka, Milwaukee, WI, USA). The first six sections containing the hippocampus bilaterally (generally from Bregma -1.3 mm to -2.5 mm) were examined for degenerating neurons using a Revolve R3 fluorescent microscope with a FITC filter (ECHO, San Diego, CA). Total number of FJC positive cells were counted in the CA1, CA3 and dentate gyrus regions of hippocampus ^39–43^ (see figure 2 C-F), degenerating neurons were counted at 20× magnification by two blinded reviewers.

### Statistical analysis

Data are presented as mean ± standard error of the mean (SEM). Statistical analyses were performed using the Real Statistics Resource Pack add-in (Release 7.6, Copyright (2013 – 2021) Charles Zaiontz. www.real-statistics.com) within Microsoft Excel. For all tests, alpha p-values less than 0.05 were considered significant. Between-group comparisons occurred via independent samples t-test, Chi-square test, one-way Analysis of Variance (ANOVA) with Tukey’s post hoc testing, or two-way ANOVA with regression, Dunn-Šidák correction, and Tukey’s post hoc testing as appropriate. The log-rank test was used for Kaplan-Meier survival curves of males and females after injury^44^.

## Supporting information

We also grouped mice by displacement into mild (0.0-2.0 mm), moderate (2.3-2.5 mm), and severe (2.7-3.0) cohorts and directly compared values to strat

Baseline motor function was assessed via rotarod in 135 male and 144 female uninjured mice. Although female mice had a greater tendency to reach the 3

Experimental research on vertebrates complied with institutional, national, and international guidelines. All procedures were approved by Duke University’s Institutional Animal Care and Use Committee. Duke University maintains an animal program that is registered and compliant with the USDA, and accredited with AAALAC, International. Recommended ARRIVE guidelines were used in design and reporting of all animal research.

The datasets generated and analyzed during the current study are not publicly available due to research funded by a private corporation and material data agreement between the university and the funding grant, but are available from the corresponding author on reasonable request.

## Authorship confirmation/contribution statement

B.M. was a project administration, writing and analysis of data and research. E.L. contributed to research methodology, investigation, data analysis, and writing of the manuscript. E.A. performed investigation and manuscript writing. B.M. was a project administration, writing and analysis of data. E.C. performed experimental investigation, data validation and review. VC performed data Investigation, and writing review. RG contributed to experimental investigation and writing review. TF performed experimental investigation and analysis, writing review & editing; BM was a project administration, performed experimental investigation, writing, review & editing of manuscript, also performed data curation, and formal analysis of data. DL contributed to methodology design, supervision of investigation, and review and editing of manuscript. BK conceptualization the study, was project administration, supervision, precured funding and supervised writing, review and editing of manuscript.

## Funding statement

Funding for this work was provided by Corticare Incorporated under a scientific research agreement.

**Supplemental Figure 1** Comparison of baseline performance on the rotarod motor task. While female mice were more likely to run the full 300 seconds at baseline, the mean latency to stopping on the task was not different between male and female C57Bl/6J mice (p > 0.05, T-test). A total of 279 mice are represented, 135 males and 144 females.

