## Supplementary material for "Optimization of a Translational Murine Model of Closed-head Traumatic Brain Injury": We also grouped mice by displacement into mild (0.0-2.0 mm), moderate (2.3-2.5 mm), and severe (2.7-3.0) cohorts and directly compared values to strat

|  | **Female Mice Rotarod** | | | | | | | | | | | |
| --- | --- | --- | --- | --- | --- | --- | --- | --- | --- | --- | --- | --- |
|  | Baseline | | | Day 1 | | | Day 3 | | | Day 6 | | |
|  | average | STD | p=value | average | STD | p=value | average | STD | p=value | average | STD | p=value |
| Mild displacement  (0-2mm) | 270.4 | 34.3 | > 0.05 | 270.4 | 34.3 | < 0.001 | 235.5 | 91.8 | < 0.05 | 260.6 | 54.0 | < 0.05 |
| Mild rotarod time  (201-300 sec) | 277.2 | 32.6 |  | 265.1 | 33.1 |  | 277.2 | 28.2 |  | 280.4 | 28.6 |  |
| Moderate displacement (2.3-2.5mm) | 266.8 | 40.4 | > 0.05 | 114.5 | 92.7 | > 0.05 | 198.3 | 85.2 | > 0.05 | 236.7 | 80.1 | > 0.05 |
| Moderate rotarod time (101-200 sec) | 261.3 | 39.3 |  | 156.4 | 24.4 |  | 230.9 | 55.0 |  | 260.1 | 41.3 |  |
| Severe displacement  (2.7-3.0 mm) | 265.4 | 37.1 | > 0.05 | 51.3 | 71.6 | < 0.05 | 126.2 | 99.3 | > 0.05 | 189.6 | 97.3 | > 0.05 |
| Severe rotarod time  (1-100 sec) | 260.8 | 39.3 |  | 29.3 | 24.2 |  | 101.8 | 84.9 |  | 169.9 | 94.0 |  |
|  | **Male Mice Rotarod** | | | | | | | | | | | |
|  | Baseline | | | Day 1 | | | Day 3 | | | Day 6 | | |
|  | average | STD | p=value | average | STD | p=value | average | STD | p=value | average | STD | p=value |
| Mild displacement  (0-2mm) | 264.8 | 33.7 | > 0.05 | 218.7 | 83.1 | < 0.05 | 252.0 | 71.9 | > 0.05 | 249.1 | 77.5 | > 0.05 |
| Mild rotarod time  (201-300 sec) | 265.7 | 34.0 |  | 266.0 | 30.2 |  | 279.8 | 28.4 |  | 279.0 | 29.0 |  |
| Moderate displacement (2.3-2.5mm) | 269.0 | 29.6 | > 0.05 | 141.2 | 87.8 | > 0.05 | 211.0 | 76.4 | > 0.05 | 246.6 | 59.1 | > 0.05 |
| Moderate rotarod time (101-200 sec) | 271.1 | 28.4 |  | 148.5 | 27.8 |  | 228.6 | 54.2 |  | 252.6 | 55.6 |  |
| Severe displacement  (2.7-3.0 mm) | 265.4 | 37.1 | > 0.05 | 51.3 | 71.6 | < 0.001 | 126.2 | 99.3 | < 0.01 | 189.6 | 97.3 | >0.05 |
| Severe RR time  (1-100 sec) | 267.2 | 30.6 |  | 44.1 | 28.1 |  | 133.0 | 79.6 |  | 187.9 | 90.4 |  |

**Supplemental data Table 1**: Closed head injury with a pneumatic piston impactor model has variability of injury between animals. To address this, we used Day one Rotarod as our assessment of injury. Rotarod performance was placed into 3 categories based on day 1 RR time; mild (201-300 seconds), moderate (100-200 seconds) and severe (<100 seconds) compared to displacement mild (0.0-2.0 mm), moderate (2.3-2.5 mm), and severe (2.7-3.0 mm) categories. All 279 mice, 135 males and 144 females were anylized separately after placing in appropriate category, a two-way ANOVA with regression was performed between each new category and that analyzing physical damage is statically different that by displacement. ANOVA for Females: Mild, F = 30.5, p < 0.001; Moderate, F= 12.2, p < 0.001; Severe, F= 7.0, p < 0.01, and ADOVA for Males: Mild, F = 21.0, p < 0.001; Moderate, F= 2.5 p > 0.05; Severe, F= 22.5, p < 0.001, with only Males at Moderate showing no significance. Post-Hock analysis was performed using one-way ANOVA using Bonferroni Alpha correction in Pairwise Mann-Whitney Test on each time point p value is displayed with in the chart along with the average rotarod time and standard deviation (STD) at each time point. The standard deviation, or variability, for mild, moderate, and severe of day 1 rotarod post injury was significantly less than using displacements (mild, moderate, severe) for both male and female animals cross the data set (p < 0.001, t-test) and confirmed by the ANOVA. Over all with every RR time, standard deviation (variability) was less when using mild, moderate, and sever RR day one times (18/18) after injury, instead of the displacement. This comparison decreasing overall variability of RR assessment of injury by approximately 50%.
