## Supplementary figures and images for "Optimization of a Translational Murine Model of Closed-head Traumatic Brain Injury"

### Baseline motor function was assessed via rotarod in 135 male and 144 female uninjured mice. Although female mice had a greater tendency to reach the 3

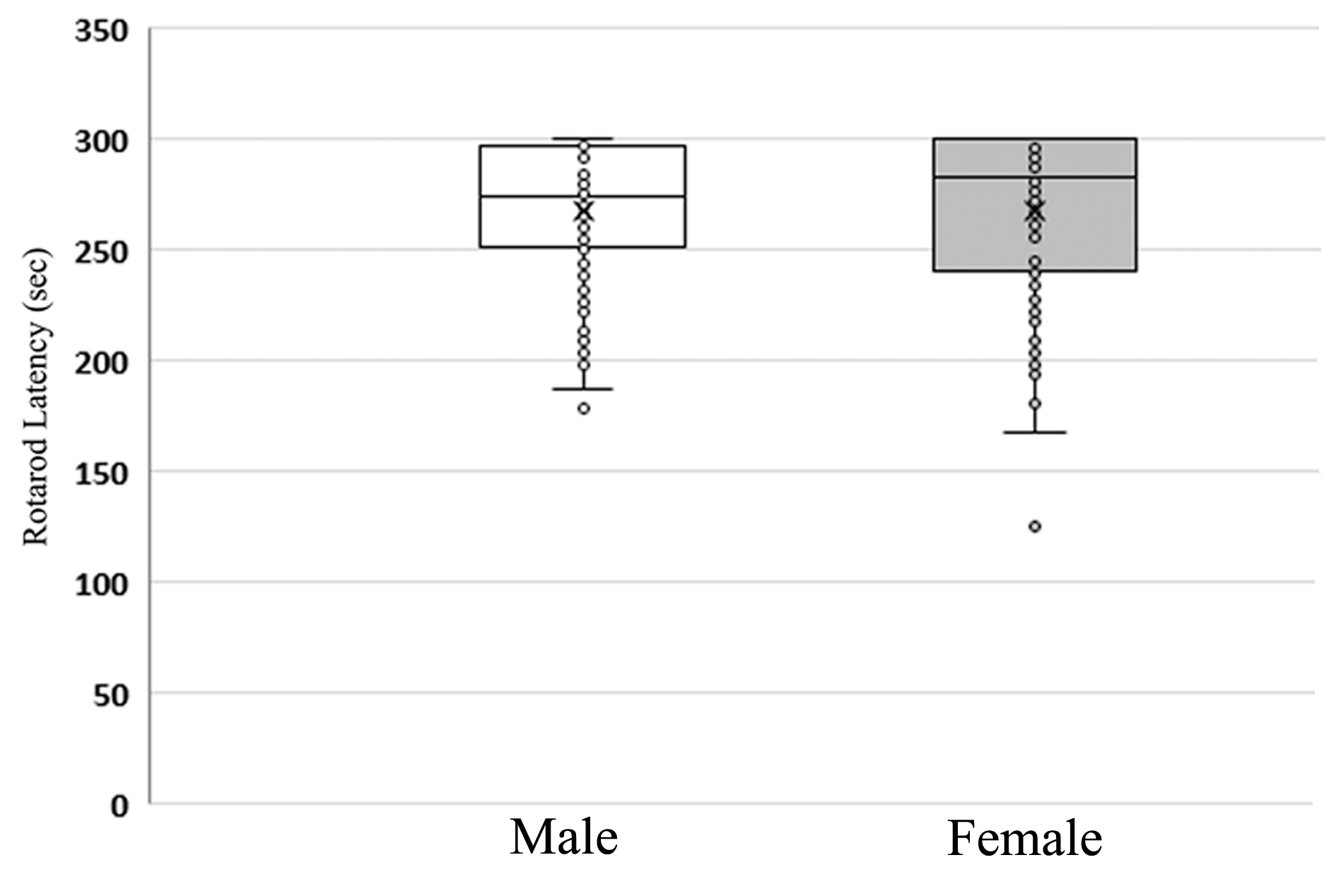
